# Pore-Resolved High-Throughput Quantification of Amyloid Aggregation and Amplification

**DOI:** 10.64898/2026.06.18.733064

**Authors:** Seokbeom Roh, Sechan Han, Minwoo Bae, Eunji Song, Taeha Lee, Dain Kang, Da Yeon Cheong, Hyungbeen Lee, Soochi Kim, Gyudo Lee

## Abstract

Amyloid fibrils are implicated in a wide spectrum of neurodegenerative and systemic disorders, yet their biological consequences are governed not only by total fibril content but also by how fibrillar species are organized, clustered, and amplified within heterogeneous populations. Conventional thioflavin T (ThT)-based assays provide sensitive sample-level readouts of β-sheet-rich material but offer limited access to the local population structure underlying amyloid aggregation and amplification. Here, we introduce the Amyloid Pore Quantification (APQ) chip, a pore-resolved geometric partitioning platform that converts heterogeneous amyloid assembly states into fluorescence intensity distributions across thousands of defined pore-level units. Using hen egg-white lysozyme as a model amyloid-forming protein, we combine length-controlled truncated amyloid nanofibrils with vacuum-assisted ThT infiltration to establish a reproducible pore fluorescence intensity reference. APQ provided quantitative concentration-dependent calibration across a 50-fold concentration range, enabling bulk-comparable quantification while preserving pore-level distribution information. Deviations from this reference resolved pH-dependent fibril clustering near the isoelectric point as pore-signal suppression and accessibility loss, captured pepsin-associated fibrillar amplification as a population-wide increase in pore intensity, and distinguished monomer- and oligomer-driven fibril processing through coupled amplification–aggregation fingerprints. When benchmarked against ensemble fluorescence measurements and AFM morphological analysis, APQ revealed assembly-state changes that were not fully represented by sample-level ThT intensity alone. These results establish APQ as a high-throughput, distribution-aware analytical framework for translating amyloid aggregation and amplification into quantitative pore-resolved fingerprints.

---

Amyloid fibrils are highly ordered, β-sheet–rich protein assemblies implicated in several neurodegenerative and systemic disorders, including Alzheimer’s disease, Parkinson’s disease, type 2 diabetes, and systemic amyloidosis.^1-5^ Although fibril deposition is a pathological hallmark, amyloid-associated toxicity and disease progression may be influenced by additional assemblies, including prefibrillar oligomers, fragmented fibrils, clustered aggregates, and seed-competent intermediate states.^6, 7^ Importantly, deposited fibrils are not necessarily inert end products; fibril surfaces and fibril fragments can catalyze secondary nucleation, support elongation, and propagate new amyloid species through seed-dependent amplification.^8-10^ Amyloid assemblies are also functional building blocks in microbial biofilms, peptide hormone storage systems, and engineered biomaterials.^11-13^ Their biological impact and material outcomes are influenced by the amount of fibrillar material present as well as the organization, fragmentation, clustering, and amplification of fibrillar species within heterogeneous populations. Quantifying amyloid assembly therefore requires analytical strategies that capture population-level organization in addition to total fibril content.

Conventional amyloid assays have been considerably valuable for monitoring fibril formation, particularly by using thioflavin T (ThT) fluorescence to detect the accumulation of β-sheet–rich amyloid structures with high sensitivity and experimental simplicity.^14^ However, fluorescence and related bulk spectroscopic measurements provide sample-level signals for fibrillar material.^15-18^ Therefore, samples with similar total β-sheet content may have comparable bulk signals, even when their underlying assembly states substantially differ in fibril length, clustering, pore accessibility, or seeding activity. This distinction is central to amyloid propagation; dispersed fibril fragments, elongated fibrils, and clustered deposits with similar total β-sheet content can differ in accessible surface area, fibril-end availability, and capacity to drive secondary amplification. These considerations point to an analytical need that is not simply greater sensitivity, but a readout that captures how ThT-positive fibrillar material is distributed across a population.

Existing amyloid characterization methods provide important but partial access to this information. Single-particle techniques such as atomic force microscopy, transmission electron microscopy, single-molecule fluorescence, and nanopore resistive-pulse sensing can resolve fibril or oligomer morphology, length, size, shape, and heterogeneity at the level of individual assemblies.^19-23^ Liquid-phase AFM analysis of cerebrospinal fluid has demonstrated that fibril length and morphology can be used for disease-stage and biomarker-related information, indicating the biological value of morphology-based amyloid analysis in clinically relevant samples, while recent AI-enabled AFM workflows further emphasize ongoing efforts to improve the reproducibility and scalability of nanoscale morphological analysis.^24, 25^ These methods are essential for structural validation, but they remain relatively labor-intensive and are not easily adapted to routine screening for many biochemical conditions. In contrast, plate-based fluorescence and amplification assays are simple, scalable, and high-throughput measurements for monitoring fibril formation and seeding activity;^26, 27^ however, they are not designed to retain unit-level population information. Therefore, a method that combines the throughput and operational simplicity of plate-based assays with unit-level distribution information is needed.

A pore-resolved geometric framework provides a route to this type of measurement. Amyloid aggregation and amplification are biochemical processes that influence how fibrillar materials occupy space. Dispersed short fibrils can enter and populate defined pore units relatively uniformly, whereas fibril-fibril clustering can generate larger assemblies that become partially or fully excluded from the pore geometry. Conversely, seed-dependent elongation or secondary processing can increase the local density of ThT–positive fibrillar materials within accessible pores. In this view, each pore-level signal is not merely a miniaturized fluorescence measurement. Instead, the distribution of signals across many geometrically defined units encodes the balance between pore-accessible fibrillar material, pore-incompatible aggregates, and amplified fibrillar populations.

Implementing this framework requires control over two conditions. First, the starting fibril population must be sufficiently uniform, because amyloid fibril contour length and length distribution can influence higher-order organization; therefore, the pore-to-pore variation should reflect assembly state changes rather than uncontrolled upstream polydispersity.^28^ Length-controlled truncated amyloid nanofibrils (t-ANFs), generated by top-down processing of preformed amyloid fibrils, provide such a reference population and allow fibrillar dimensions to be matched to the analytical geometry of the pore array.^8, 29^ Second, the staining and loading processes must consistently deliver ThT–positive fibrillar material into pore units to ensure the measured signal reflects fibril accessibility and local fibrillar density rather than infiltration variability.^30^ Although miniaturized amyloid assays were used to improve the speed and throughput of amyloid detection,^31-33^ assay-style platforms that combine operational simplicity with pore-based population fingerprinting remain underdeveloped. When length-controlled fibrils and reproducible infiltration are combined, uniformly loaded t-ANFs can be used to establish a stable pore fluorescence intensity reference, whereas deviations from this reference may report assembly state-dependent changes.

Here, we introduce the Amyloid Pore Quantification (APQ) chip, a pore-resolved analytical platform that converts heterogeneous amyloid assembly states into fluorescence intensity distributions across thousands of geometrically defined pore-level units. Using hen egg-white lysozyme (HEWL) as a model amyloid-forming protein,^34^ we combined length-controlled t-ANFs with vacuum-assisted ThT infiltration to establish a reproducible pore intensity baseline and detect assembly state-dependent deviations from this reference state. APQ was used for three amyloid processes: pH-dependent fibril clustering near the isoelectric point (pI), protease-associated fibril amplification, and monomer- or oligomer-driven fibril processing. By comparing APQ readouts with ensemble fluorescence measurements, we show that the platform retains compatibility with conventional amyloid quantification while adding distribution-resolved information on pore accessibility, local fibrillar density, and aggregation-dependent exclusion. Thus, APQ may be a distribution-based analytical framework for converting amyloid aggregation and amplification into high-throughput pore-based fingerprints.

## RESULTS AND DISCUSSION

### Design of the APQ Chip and Uniform t-ANFs

The APQ chip is built around a layered PDMS–adhesive–PDMS architecture that sandwiches a commercial track-etched polycarbonate membrane filter as the analytical core (Figure 1a). Two PDMS slabs (20 × 20 mm²), each carrying a central 10 mm circular opening, define the analysis window, while a 70 μm-thick adhesive layer secures the membrane and prevents sample bypass around its edges. The exposed membrane region presents an array of cylindrical 10 μm pores that act as parallel analytical compartments; the inset of Figure 1a highlights the dense, evenly distributed pore field available for pore-resolved readouts. Statistical analysis of the membrane confirmed a narrow pore-diameter distribution centered at 11.49 ± 0.22 μm, with a small, well-separated minor population corresponding to angled or merged multi-pores (Figure 1b), providing a stable geometric baseline against which fibril-dependent fluorescence variations can be reliably interpreted. Given the nominal pore diameter of 10 μm and a membrane thickness of 16 μm, each pore defines a cylindrical compartment of approximately 1.3 pL (Figure 1c), placing the APQ chip in the picoliter-partition regime relevant to membrane-based digital assays.

**Figure 1.**
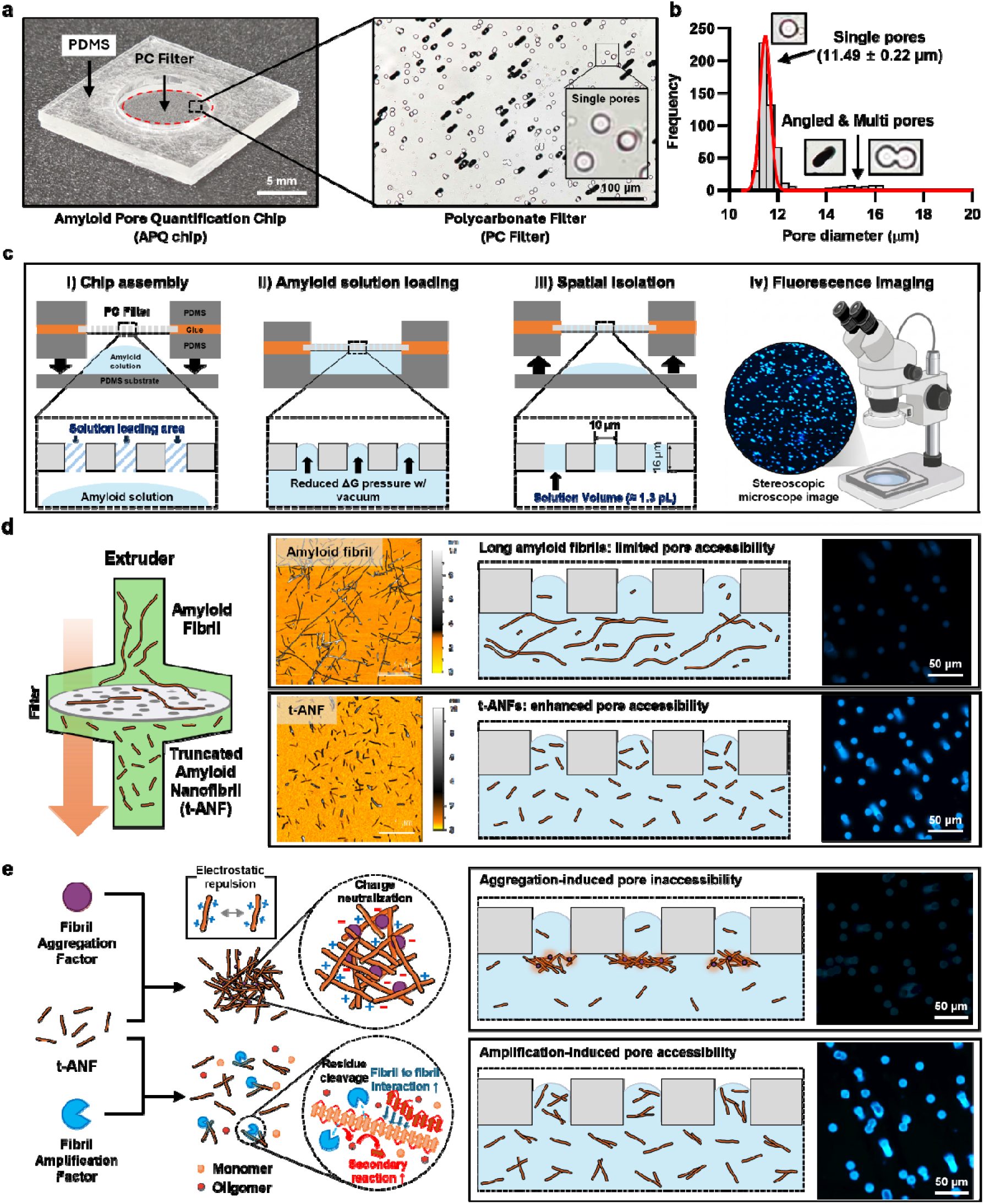
Concept and design of the APQ platform for pore-resolved amyloid quantification. (a) Image and schematic of the APQ chip, composed of a PDMS–adhesive–PDMS structure incorporating a track-etche polycarbonate membrane filter; the inset shows the exposed pore field. (b) Membrane pore diameter distribution. (c) Workflow of APQ chip loading, vacuum-assisted pore isolation, and fluorescence imaging, with the estimated single-pore volume. (d) Generation of t-ANFs by extrusion-based mechanical processing and comparison with conventional long fibrils using AFM, schematic illustrations, and pore-level fluorescence imaging. (e) Conceptual framework for aggregation-associated pore exclusion and amplification-associated pore enrichment.

To match the pore geometry to the fibril population being measured, we generated t-ANFs by extruder-based top-down processing (Figure 1d). AFM, schematic, and fluorescence comparisons (Figure 1d) show that t-ANFs distribute uniformly across pores, whereas conventional long fibrils give sparse, heterogeneous pore loading. This t-ANF–induced enhancement of pore accessibility was quantitatively confirmed using single-pore intensity tracking across N ≈ 600 pores at 0.1 wt% (w/v; 1 mg/mL); t-ANF samples produced consistently brighter pores (∼13,000 a.u.) than long-fibril samples (∼2,000 a.u.), corresponding to an approximately six-fold difference at identical bulk fibril content (Figure S1). This separation was further supported by a Z-factor of 0.614, confirming a robust pore-level assay window between conventional long fibrils and pore-accessible t-ANFs. The improved length uniformity translates directly into more reproducible pore-level fluorescence readouts and allows individual pores to be treated as quantitative analytical units. Building on the combination of a regular pore array and a uniform fibril substrate, we defined the analytical framework of the APQ chip in terms of two complementary pore-level signatures: aggregation-associated pore exclusion and amplification-associated pore enrichment (Figure 1e). Aggregation-associated pore exclusion captures pore-level signal loss caused by fibril–fibril clustering and pore incompatibility, whereas amplification-associated pore enrichment captures signal increases arising from seed-dependent growth or secondary processing.

### Optimization of Vacuum-Assisted ThT Staining

We next optimized vacuum-assisted ThT staining so that pore-level fluorescence faithfully reports fibril content rather than infiltration variability (Figure 2). The APQ chip was placed in a vacuum desiccator and evacuated to −0.85 bar gauge pressure (Figure 2a), creating a pressure differential that drives the t-ANF/ThT solution into individual pores. Time-lapse imaging (Figure 2b) revealed a progressive build-up of pore fluorescence between 0 and 10 min, and the corresponding pore-resolved heatmap (Figure 2c) showed that pore-to-pore signal converged to a saturated, uniform pattern by ∼5–10 min (12.54 ± 0.93 a.u. at 5 min and 12.11 ± 1.32 a.u. at 10 min, versus 2.95 ± 0.30 a.u. at 0 min). We therefore selected 10 min as the standard infiltration time, a condition that yields complete pore filling with consistent inter-pore fluorescence and minimal residual variability.

**Figure 2.**
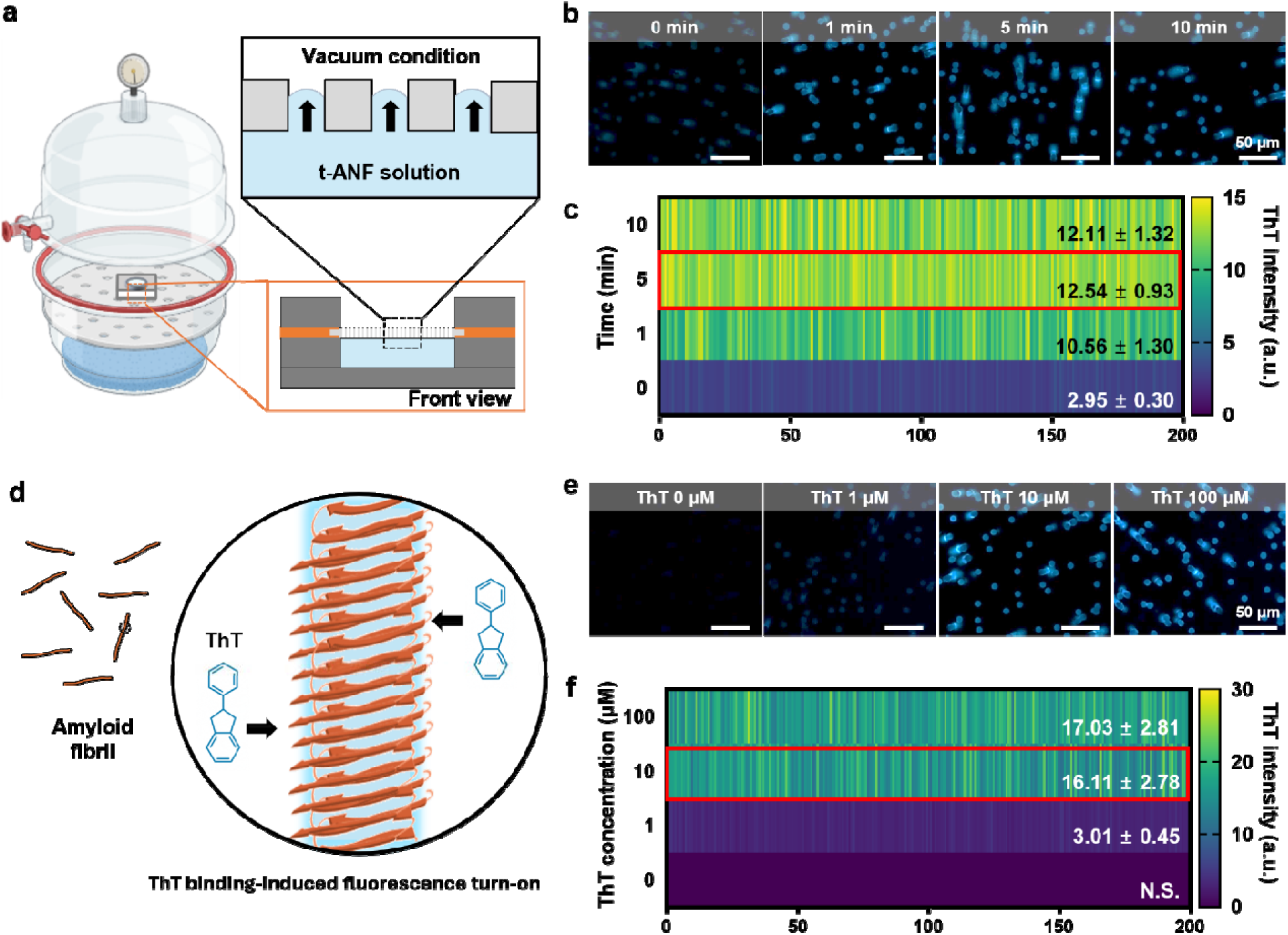
Optimization of vacuum-assisted ThT staining within the APQ platform. (a) Schematic of the vacuum-driven infiltration setup. The APQ chip was placed in a vacuum desiccator and evacuated to −0.85 bar gaug pressure, allowing the t-ANF–ThT solution to be drawn into individual pores. (b) Time-lapse fluorescence images of ThT–stained pores acquired at 0, 1, 5, and 10 min during vacuum infiltration. (c) Heatmap of pore-resolved ThT intensity as a function of infiltration time, showing the temporal evolution of pore filling and fluorescence signal. (d) Molecular schematic illustrating ThT binding to the cross-β architecture of amyloid fibrils. (e) Fluorescence images of t-ANF-filled pores stained at varying ThT concentrations (0, 1, 10, and 100 μM). (f) Heatmap of pore-resolved ThT intensity as a function of ThT concentration.

The ThT concentration was then optimized in light of the molecular basis of ThT–fibril recognition (Figure 2d): ThT inserts into the cross-β architecture of amyloid fibrils, and its quantum yield rises sharply upon binding, making pore-level fluorescence an operational reporter of ThT-accessible fibrillar material within each pore.^35^ Fluorescence images (Figure 2e) and the corresponding pore-resolved heatmap (Figure 2f) acquired at 0, 1, 10, and 100 μM ThT showed that 10 μM provided strong, well-resolved per-pore signal (16.11 ± 2.78 a.u.) with low background. In contrast, 100 μM only marginally increased the mean intensity (17.03 ± 2.81 a.u.) without adding analytical contrast and with higher background. We therefore used 10-min vacuum infiltration with 10 μM ThT throughout the remainder of this study—conditions that combine reproducible pore filling with a saturation-limited ThT response across the pore array.

### Quantitative Calibration across t-ANF Concentrations

To establish the APQ chip as a quantitative platform, we calibrated pore-level fluorescence against t-ANF concentration (Figure 3). Fluorescence images of the pore array loaded with 0.01, 0.05, 0.1, and 0.5 wt% t-ANF showed a clear, monotonic brightening of individual pores with increasing fibril concentration (Figure 3a). Magnified single-pore views with line-scan profiles (Figure 3b) confirmed that the intensity increase was localized within the pore region itself, without spurious contributions from the surrounding membrane. Aggregating these line scans across multiple pores per condition yielded a quantitative concentration-dependent gray-value profile that rises steeply within the 10 μm pore footprint and drops off sharply at the pore boundaries (Figure S2), demonstrating that the per-pore signal is genuinely confined to the pore lumen and scales reproducibly with t-ANF concentration. Across 500 pores per condition, the single-pore intensity scatter and corresponding histograms (Figure 3c) shifted toward higher values with increasing t-ANF concentration (9,278 ± 1,531; 12,877 ± 618; 22,905 ± 1,497; and 35,939 ± 3,678 a.u. for 0.01, 0.05, 0.1, and 0.5 wt%, respectively) while preserving narrow, unimodal distributions. The chip-to-chip reproducibility of this pore-level readout was further verified by comparing three independent APQ chips loaded with the same 0.1 wt% t-ANF stock, which yielded comparable pore-intensity distributions across chips (Figure S3).

**Figure 3.**
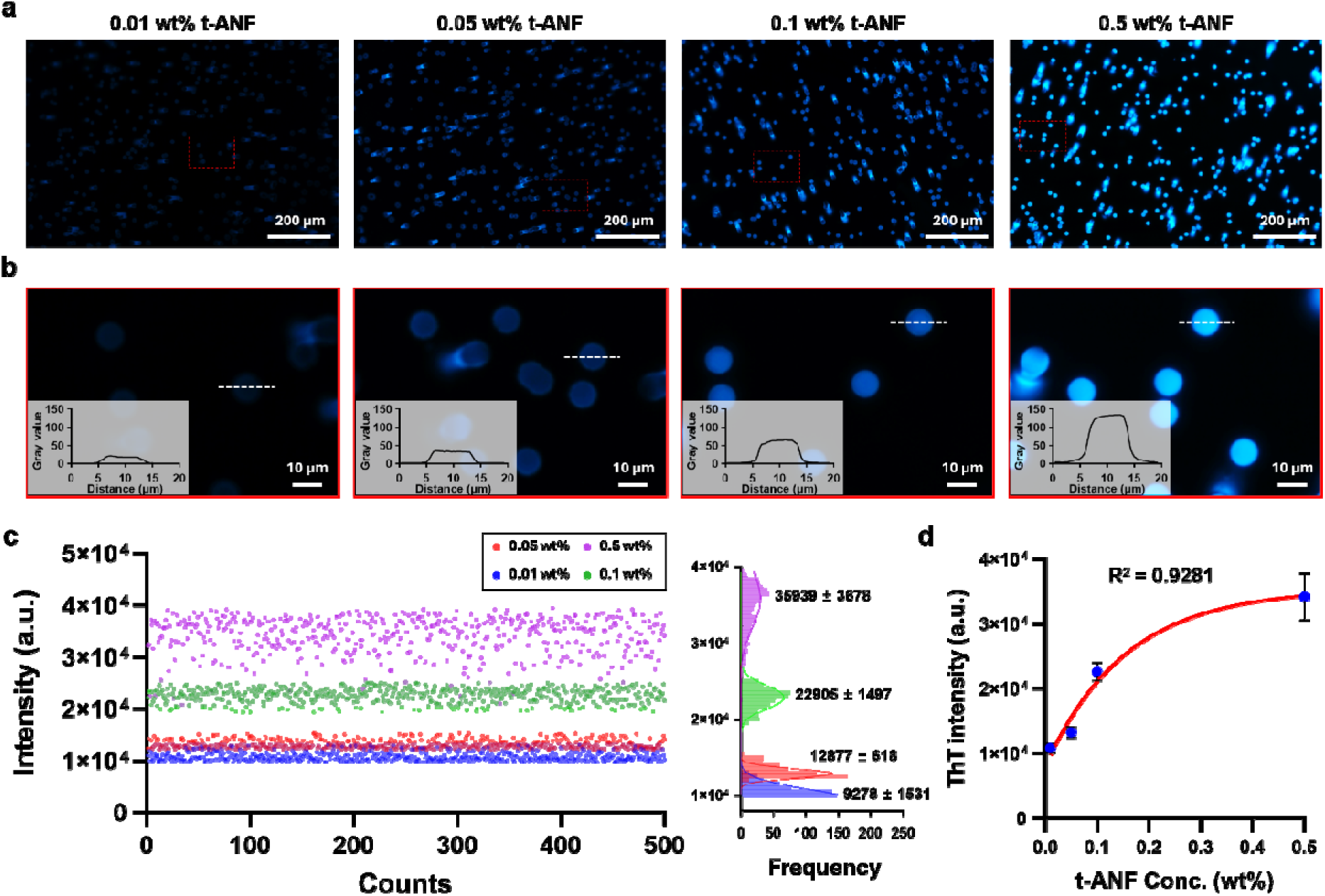
Quantitative calibration of the APQ platform across t-ANF concentrations. (a) Fluorescence images of pores loaded with t-ANF solutions at 0.01, 0.05, 0.1, and 0.5 wt%. (b) Magnified views of representative pores from each concentration, with insets showing line-scan gray-value profiles across individual pores (dashed lines). (c) Single-pore ThT intensity distributions across 500 pores (left) and the corresponding frequency histograms (right) for the four t-ANF concentrations. (d) Calibration curve of mean integrated pore fluorescence intensity versus t-ANF concentration, fitted using a nonlinear exponential-plateau model (*R*^2^ = 0.9281).

Because the t-ANF population is length-controlled and the loading/staining conditions are fixed, the pore array can be treated as an empirical partitioning geometry in which the pore-intensity distribution reports the amount and accessibility of fibrillar material. As a conceptual approximation, if effective fibrillar units enter pores independently with an average occupancy *λ*, the probability of observing *m* effective units in a pore follows:

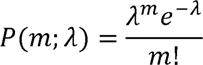

This expression was used only as a statistical rationale for uniform pore partitioning and was not fitted to the fluorescence data. In the present high-occupancy regime, each pore contains many fibrillar units; therefore, the relevant experimental signature is the preservation of a narrow, unimodal pore-intensity distribution whose mean shifts with fibril loading.

The mean integrated pore intensity, 〈*I*_total_〉, increased monotonically with t-ANF concentration, *c*_t-ANF_, and was fitted using an empirical nonlinear exponential-plateau model:

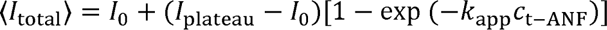

where *I*_0_ is the fitted baseline intensity, *I*_plateau_ is the saturation-limited plateau intensity, and *k*_app_ is the apparent concentration-response coefficient. The fit yielded *R*^2^ = 0.9281 across the tested 50-fold t-ANF concentration range.

### pH-Dependent Amyloid Aggregation

We first applied the calibrated APQ chip to map pH-dependent aggregation of HEWL t-ANFs (Figure 4). Identical t-ANF stocks were adjusted to pH 9, 10, 11, and 12 prior to loading. Fluorescence image (Figure 4a) and pore-resolved heatmaps (Figure 4b) revealed clear pH-dependent modulation of pore-level signal, with the most pronounced change occurring at pH 11—the condition closest to the isoelectric point (pI) of HEWL (pI ≈ 10.7–11.3; Figure 4d, inset)—where the mean pore intensity dropped to 681 ± 61 a.u., compared to 1,765 ± 259, 1,247 ± 181, and 1,713 ± 255 a.u. at pH 9, 10, and 12, respectively. Single-pore intensity scatter plots (Figure 4c) and frequency distributions (Figure 4d) further showed that this change is not a simple bulk shift: at pH 11, the distribution narrowed sharply (693 ± 68 a.u.) and shifted to lower intensities, while at pH 9 and 12 the distributions remained broader and centered at substantially higher pore intensities (1,719 ± 299 and 1,724 ± 312 a.u., respectively). The departure from the narrow, unimodal reference distribution established under uniform t-ANF loading (Figure 3c) thus serves as a direct, distribution-level signature of aggregation, in which the pI-centered loss of pore signal—rather than a simple shift in the mean—reports on enhanced fibril–fibril association outside the pore-accessible size window. Closer inspection of the pH 10 distribution further revealed an instructive sub-population: although the majority of pH 10 pores clustered around 1,247 ± 181 a.u., approximately 6.3% (∼63 of *N* = 1000) fell within the low-intensity regime characteristic of pH 11 (Figure 4c,d), with a sub-population mean of ∼693 ± 68 a.u.—essentially indistinguishable from the pH 11 main population. We interpret this minority population as reflecting heterogeneous aggregation within the pH 10 sample, possibly arising from local variations in ionic/protonation environment or fibril–fibril contact history that generate pI-like aggregation behavior in a subset of pore-accessible populations.^36-38^ Such minority populations are readily masked by ensemble averaging and illustrate the type of heterogeneity that distribution-resolved measurements can capture. Consistent with this interpretation, parallel bulk ThT measurements of the same pH-adjusted t-ANF samples showed only minor, statistically non-significant differences across pH 9–12 (Figure S4), whereas APQ resolved a pronounced pore-signal suppression at pH 11. This contrast illustrates the central analytical advantage of APQ: pore-resolved readouts convert aggregation-dependent accessibility changes into quantitative distribution-level fingerprints. Because bulk ThT integrates fluorescence over the sample volume, it does not directly resolve how ThT-positive material is partitioned among dispersed fibrils, clustered aggregates, and pore-incompatible assemblies. In other words, the cross-β signature of an individual t-ANF and of a clustered t-ANF super-aggregate is essentially the same (Figure S4). The pronounced ∼3-fold pore-signal drop that APQ resolves at pH 11 (Figure 4b) therefore represents an aggregation-dependent accessibility change that is not directly resolved by bulk ThT fluorescence.

**Figure 4.**
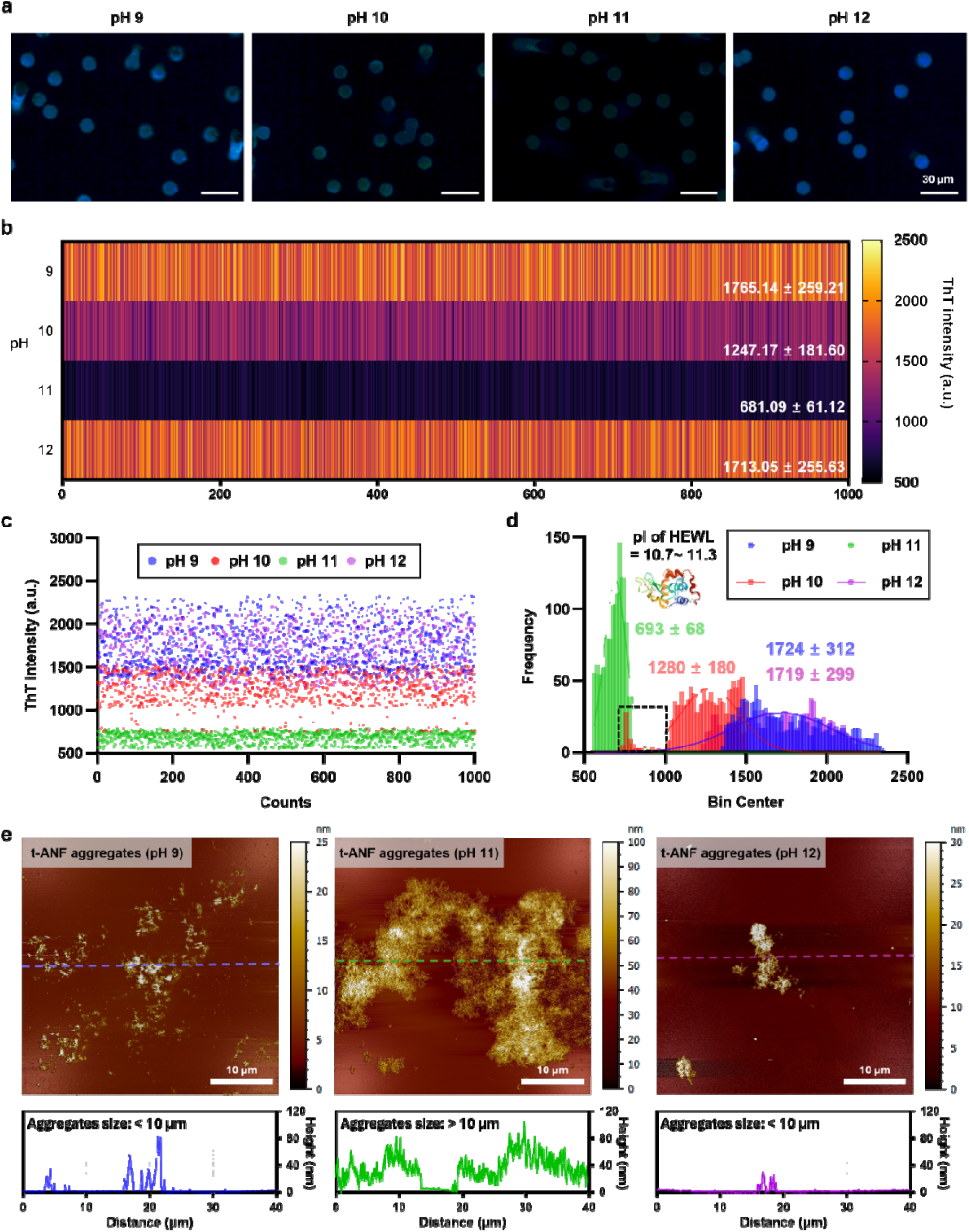
APQ analysis of pH-dependent amyloid aggregation. (a) Fluorescence images of t-ANF-loaded pores at pH 9, 10, 11, and 12. (b) Pore-resolved ThT intensity heatmap and (c) scatter plot for each pH condition. (d) Frequency distributions of pore ThT intensities, annotated with the HEWL pI range; the dashed box marks low-intensity pores attributed to aggregate exclusion, and the inset shows the HEWL structure (PDB ID: 8RUS.^42^). (e) AFM topography images of t-ANF aggregates at pH 9, 11, and 12.

AFM topography of the corresponding aggregates (Figure 4e) supported this size-exclusion interpretation. At pH 9, t-ANFs assembled into small (<10 μm), discrete clusters of short fibrillar species; at pH 11, near pI, the same nanofibrils formed denser, interconnected superstructures with footprints exceeding 10 μm—beyond the pore diameter—explaining the strongly suppressed pore-level signal at this condition; at pH 12, smaller (<10 μm) aggregates re-formed, consistent with the recovery of pore-level fluorescence. The pore-resolved fluorescence distributions and AFM morphologies thus converge on a coherent picture in which charge neutralization at pI drives fibril–fibril association into pore-incompatible super-aggregates that are physically excluded from the 10 μm pores. This interpretation is consistent with prior surface-potential analyses of amyloid aggregates, in which Kelvin probe force microscopy showed that fibril surface charge passes through zero near the protein’s pI;^39^ the resulting electrostatic neutralization is what enables fibril–fibril association and drives the assembly of clustered, interconnected superstructures. Comparable charge-mediated assembly behavior has been reported for HEWL fibrils interacting with oppositely charged polyelectrolytes, where electrostatic complexation drives fibril clustering and three-dimensional network formation.^40^ Statistical analysis of single-fibril AFM images, in particular, has been established as a powerful approach for characterizing such heterogeneous fibrillar populations,^20, 41^ and the pore-resolved fluorescence distributions reported here provide a complementary, throughput-oriented readout that captures the same heterogeneity at the level of thousands of pores. The APQ chip thus translates a continuous, charge-driven aggregation behavior into a distribution-based quantitative fingerprint that is accessible from a single fluorescence acquisition, and it does so with fibril usage and instrumentation requirements that are well suited to routine screening of aggregation conditions.

### Pepsin-Induced Secondary Amplification of t-ANFs

We next examined whether the APQ chip can capture amplification—seed-dependent processes that increase or redistribute amyloid-associated signal^9^—using pepsin as a representative protease (Figure 5). t-ANFs were incubated for 3 h at 37 °C under four conditions: t-ANF alone, t-ANF + pepsin, t-ANF + pepsin + protease inhibitor, and t-ANF + heat-denatured pepsin. Fluorescence images of the pore array (Figure 5a) showed visibly higher and more uniformly bright pores under pepsin treatment, while the inhibitor co-treated samples produced an intermediate signal and the denatured-pepsin conditions converged near baseline (*i.e.*, t-ANF only).

**Figure 5.**
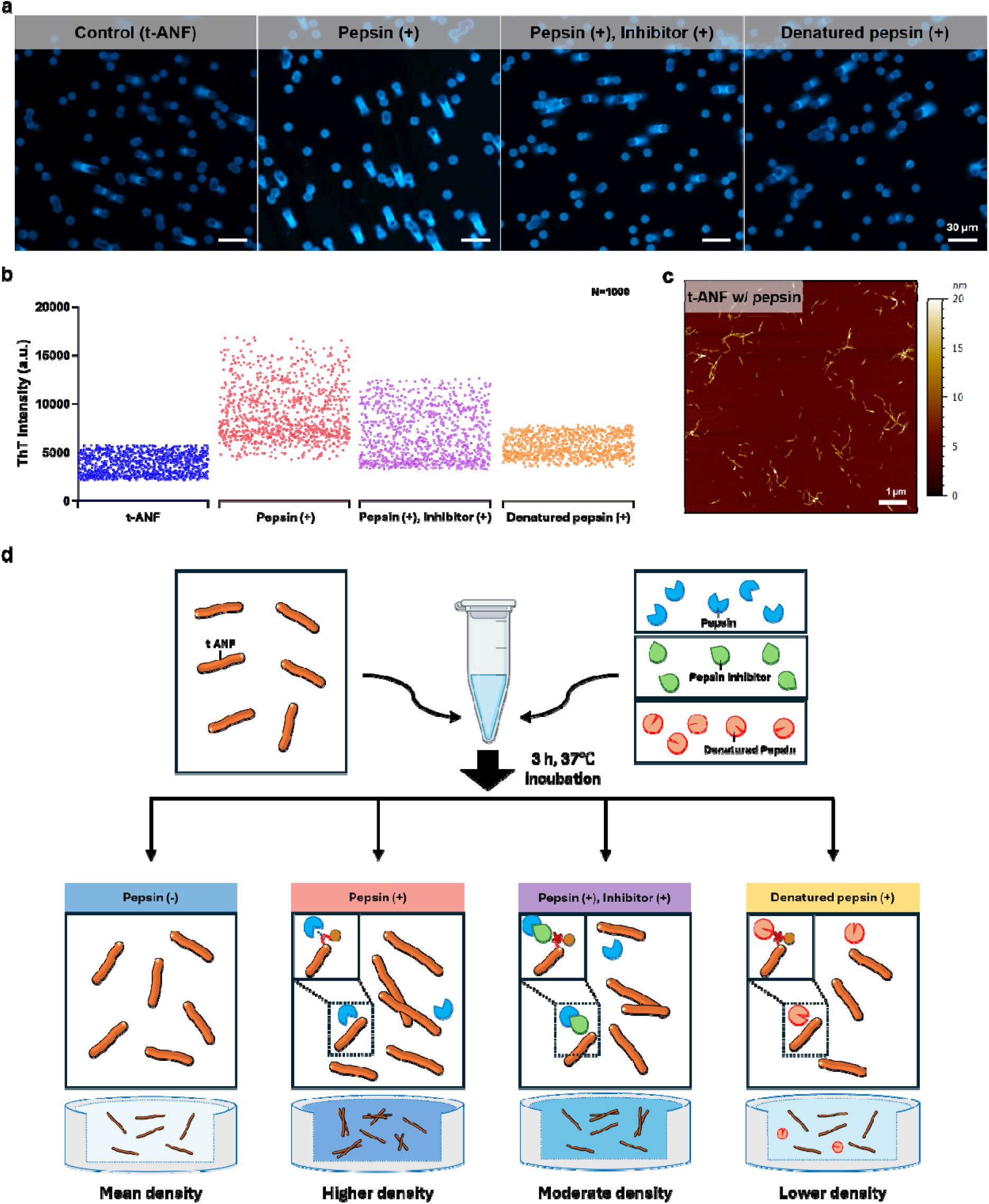
APQ analysis of pepsin-associated t-ANF amplification. (a) Representative fluorescence images of APQ pores loaded with t-ANF alone, t-ANF + pepsin, t-ANF + pepsin + protease inhibitor, and t-ANF + heat-denature pepsin. Scale bars, 30 μm. (b) Single-pore ThT intensity distributions for each condition; each dot represents on pore (*N* = 1000 pores per condition). (c) AFM topography image of pepsin-treated t-ANFs. Scale bar, 1 μm. (d) Schematic of the pepsin treatment workflow and corresponding pore-level readout after 3 h incubation at 37 °C.

Pore-level distributions from N =1000 analyzed pores per condition (Figure 5b) sharpened this picture. Pepsin treatment shifted the distribution toward markedly higher intensities, indicating an increased density of ThT-positive fibrillar material within each pore. Adding the protease inhibitor partially suppressed this shift, while heat-denatured pepsin returned the distribution close to baseline, although a small residual elevation above the t-ANF-only control was still discernible (Figure 5b). This residual signal may reflect a minor enzymatic-activity-independent contribution, potentially from weak nonspecific protein–fibril interactions. The full distribution—rather than only the mean—shifted, demonstrating that the increase in pore signal is a population-wide phenomenon rather than the consequence of a small number of outlier pores. This pore-level enhancement was mirrored by a complementary bulk-ThT measurement on the same samples, which showed a highly significant (∼24%) increase in ensemble fluorescence between t-ANF control (∼13,300 a.u.) and pepsin-treated t-ANF (∼16,500 a.u.) (Figure S5). The convergence of bulk-level and pore-level enhancement here—unlike the cases in which APQ resolved accessibility changes that were not fully reflected in bulk ThT—indicates that pepsin-induced secondary processing increases the ThT-positive fibrillar signal while preserving pore accessibility, consistent with amplification-like fibrillar processing rather than redistribution into pore-incompatible aggregates. AFM imaging of the pepsin-treated sample (Figure 5c) revealed coexisting fragmented species and newly elongated fibrillar segments, consistent with limited pepsin-mediated processing of t-ANFs producing additional fibril ends and surfaces that support downstream fibrillar assembly. Quantitative analysis of single-fibril AFM populations (Figure S6) supported this picture: while the length distributions of control and pepsin-treated samples both peaked below ∼250 nm, the pepsin-treated population developed a long-length tail extending beyond ∼2,000 nm, and the height distribution shifted from a sharp ∼1–2 nm peak in the control to a broad multimodal distribution centered around 5–10 nm with a tail extending beyond 20 nm—directly consistent with the appearance of thicker, partially elongated fibrillar species after proteolytic exposure. Comparable behavior has been reported for lysozyme-derived amyloid: among multiple amyloid systems, lysozyme amyloid exhibits the highest pepsin resistance and undergoes proteolysis-driven secondary nucleation that leads to fibril proliferation rather than degradation, accompanied by an increase in fibril thickness and stiffness.^43^

We interpret these observations as a working hypothesis (Figure 5d): pepsin may cleave exposed or flexible regions of t-ANFs, altering surface residues, charge distribution, or hydrophobic exposure in a way that enhances fibril–fibril interaction and supports secondary fibrillar growth, leading to the high pore densities observed in APQ. The graded responses—high signal under pepsin, intermediate signal under pepsin + inhibitor, and near-baseline signal under denatured pepsin—indicate that this effect predominantly tracks the enzymatic activity of pepsin, with only a minor non-enzymatic contribution from the polypeptide itself. Direct biochemical confirmation of cleavage sites and surface chemistry will be valuable in future work; here, we emphasize that the APQ chip, jointly with bulk ThT (Figure S5) and AFM-resolved morphology statistics (Figure S6), resolves this enzymatic amplification as a defined, distribution-level fingerprint that distinguishes pepsin-active, pepsin-inhibited, and pepsin-inactive states from a single fluorescence acquisition.

### Monomer- and Oligomer-Induced t-ANF Amplification and Isoelectric Aggregation

Finally, we tested whether APQ can distinguish precursor-dependent t-ANF processing and the downstream aggregation behavior of the resulting fibrillar populations (Figure 6). Three samples were prepared under matched conditions in pH 2 DW: t-ANFs alone, t-ANFs co-incubated with freshly filtered HEWL monomeric precursors, and t-ANFs co-incubated with HEWL oligomeric precursors (Figure 6a; Stage 1). All samples were incubated for 3 h at 60 °C. This temperature was selected because ThT fluorescence remains substantially retained up to ∼60 °C, whereas higher temperatures produce a sharp decrease in signal due to thermal dissociation (Figure S7).^44, 45^ After the amplification incubation, each sample was loaded directly onto the APQ chip. The same incubated samples were then adjusted to pH 10 to induce downstream aggregation under a mild near-isoelectric condition and reloaded onto fresh chips for the pH-adjusted aggregation measurement (Figure 6a; Stage 2).

**Figure 6.**
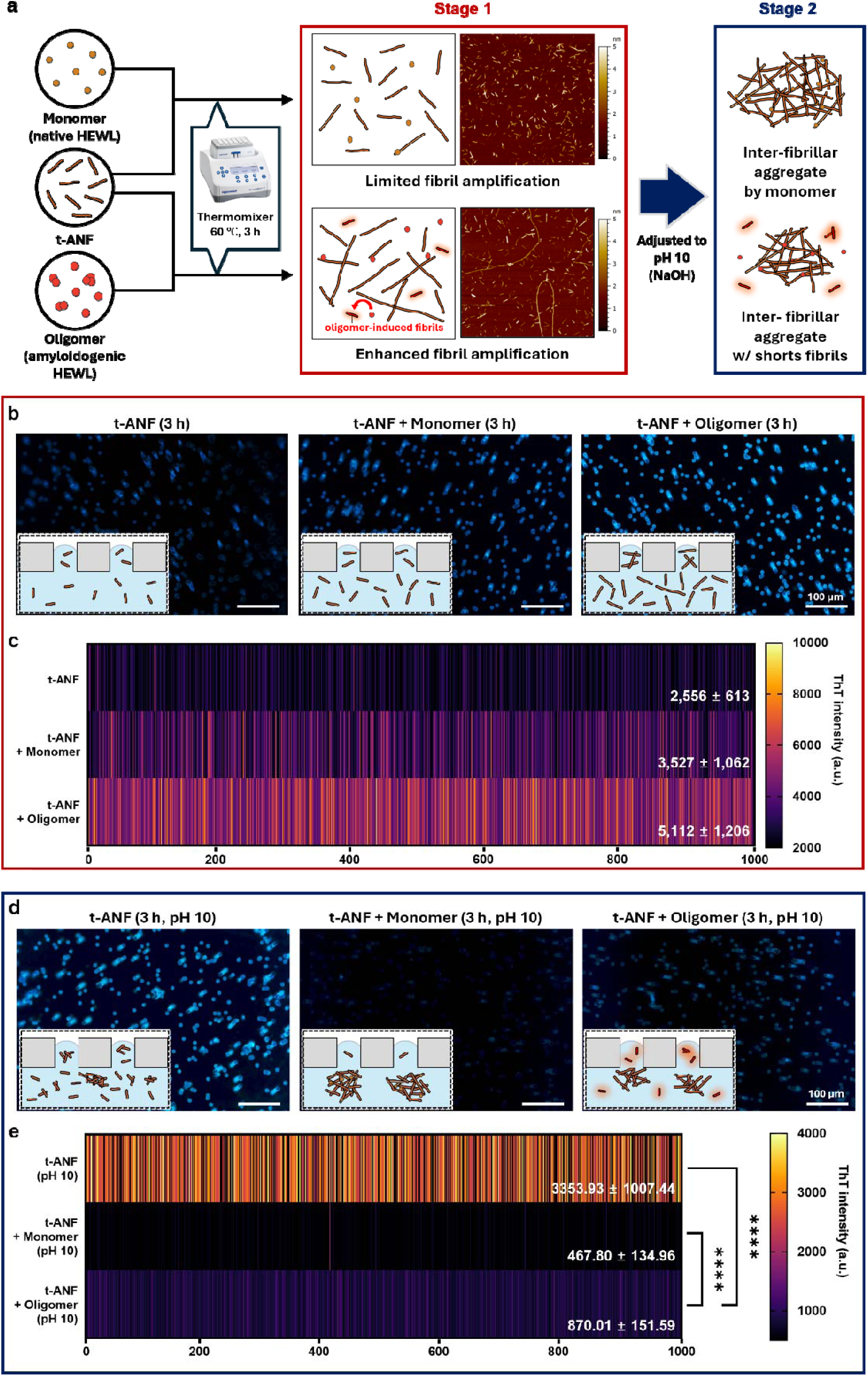
APQ-based discrimination of monomer- and oligomer-driven t-ANF amplification and downstream aggregation. (a) Schematic of the two-stage workflow. t-ANFs were incubated alone or with HEWL monomeric or oligomeric precursors at pH 2 and 60 °C for 3 h, followed by pH adjustment to 10 to induce downstream aggregation. (b) Representative APQ fluorescence images after the amplification incubation. (c) Pore-resolved ThT intensity heatmaps for the corresponding Stage 1 samples. (d) Representative APQ fluorescence images after pH 10 adjustment. (e) Pore-resolved ThT intensity heatmaps for the corresponding Stage 2 samples. *N* = 1000 pores per condition. Scale bars, 60 μm.

Fluorescence images and pore-resolved heatmaps acquired after the amplification incubation showed a clear precursor-dependent increase in pore intensity (Figure 6b,c). The t-ANF control produced the lowest mean pore intensity (2,556 ± 613 a.u.), whereas monomer co-incubation increased the signal to 3,527 ± 1,062 a.u. and oligomer co-incubation further increased it to 5,112 ± 1,206 a.u. across *N* = 1000 pores. This ordering—control < monomer < oligomer—supports a precursor-dependent difference in apparent fibrillar amplification under matched processing conditions. The oligomer-derived sample produced the strongest pore-level signal, consistent with the ability of oligomeric populations to participate in amyloid growth pathways.^46-48^ AFM analysis supported this assignment: t-ANF-only samples retained predominantly short, fragment-like morphologies (Figure S9a), whereas monomer-coincubated samples showed modest fibrillar growth and oligomer-coincubated samples developed longer and denser fibrillar networks over the same time course (Figure S9b). In contrast, bulk ThT measurements did not resolve a significant difference between monomer- and oligomer-treated t-ANFs (Figure S8), indicating that APQ preserves distribution-level information that is obscured by ensemble averaging.

Analysis after pH 10 adjustment revealed an inversion of the amplification-incubation hierarchy (Figure 6d,e). The t-ANF control retained a substantial pore-level signal (3,353 ± 1,007 a.u.), whereas the monomer- and oligomer-coincubated samples showed lower pore intensities of 467 ± 134 and 870 ± 151 a.u., respectively. Because Stage 1 APQ and AFM data indicated increased fibrillar processing in the precursor-treated samples, and bulk ThT showed comparable ensemble β-sheet signals (Figure S8), the reduced Stage 2 pore intensities are unlikely to reflect simple loss of fibrillar material. Instead, they are consistent with the formation of larger, pore-incompatible aggregates that are excluded from the 10 μm pores, following the same size-exclusion principle observed in the pH-dependent aggregation experiment (Figure 4).

AFM images of the Stage 2 samples further supported this interpretation (Figure S10). In the +monomer condition, monomer addition can promote fibril-end growth, lateral association, and fibril-catalyzed secondary nucleation on t-ANF templates.^49-51^ After pH 10 adjustment, these processed fibrillar species formed dense micrometer-scale aggregates, reducing the fraction of pore-accessible t-ANFs and producing the lowest APQ signal among the three Stage 2 conditions. In the +oligomer condition, the precursor population is expected to be more heterogeneous and structurally dynamic, potentially containing both elongation-competent and kinetically trapped species. ^52-54^ Under near-isoelectric aggregation conditions,^55^ this heterogeneity appears to produce a mixed product population containing both large aggregates and shorter fibril-like species (Figure S10). The remaining short species can still enter the 10 μm pores, explaining the residual Stage 2 signal in the oligomer condition relative to the monomer condition.

Together, the sequential APQ analysis separates precursor-dependent amplification from downstream aggregation accessibility. Stage 1 reports the apparent amplification hierarchy of the fibrillar products, whereas Stage 2 reports how those products are redistributed into pore-compatible or pore-incompatible assemblies after pH-triggered aggregation. Thus, monomer- and oligomer-driven pathways are not collapsed into a single ThT value but are resolved as distinct amplification–aggregation fingerprints across successive APQ measurements (Figure S8).

## CONCLUSIONS

We introduced the Amyloid Pore Quantification (APQ) chip, a PDMS–membrane platform that uses the pores of a commercial track-etched polycarbonate filter as parallel analytical units for high-throughput amyloid analysis. By combining length-controlled truncated amyloid nanofibrils (t-ANFs), vacuum-assisted ThT staining, and pore-level image analysis, APQ converts heterogeneous amyloid populations into fluorescence intensity distributions across thousands of geometrically defined pores. This format provides bulk-comparable quantification while retaining distribution-level information on pore accessibility and local fibrillar density.

Across the tested conditions, APQ resolved both calibration behavior and assembly-state-dependent deviations from the t-ANF reference state. Concentration calibration over a 50-fold t-ANF range produced a monotonic pore-intensity response that was captured by an empirical exponential-plateau fit (*R*^2^ = 0.9281), establishing a reproducible reference distribution for pore-accessible t-ANFs. pH-dependent measurements near the isoelectric point of HEWL revealed charge-mediated aggregation as a suppression of pore-accessible fluorescence, consistent with AFM-observed formation of pore-incompatible aggregates. Pepsin-mediated processing produced a population-wide increase in pore intensity, with graded responses across active, inhibited, and heat-denatured pepsin conditions, supporting an enzymatic contribution to amplification-like fibrillar processing. Monomer- and oligomer-driven reactions further demonstrated the ability of APQ to distinguish coupled amplification–aggregation behavior: measurements after the amplification incubation resolved a control < monomer < oligomer hierarchy, whereas measurements after pH 10 adjustment revealed a monomer < oligomer < t-ANF control ordering that reflected differences in the remaining pore-accessible fibrillar fraction.

These results position APQ as a complementary method to conventional amyloid assays. Plate-based ThT measurements provide efficient ensemble fluorescence readouts, whereas APQ adds pore-resolved distribution information from a single fluorescence acquisition. The platform is fabricated from commercially available components, requires no specialized microfabrication, and is compatible with routine laboratory imaging workflows. APQ is also conceptually complementary to capillary-flow and microfluidic amyloid analysis platforms,^31, 32, 56^ extending bulk-quantitative readouts with pore-level population information. The same pore-resolved strategy may be extendable to other amyloid-forming proteins, inhibitor screening, seeding-amplification assays, and distribution-aware analysis of complex amyloid-containing samples, including biomarker-oriented amyloid analysis.^57^ More broadly, this work demonstrates that membrane-pore architectures can be repurposed from passive filtration elements into high-density analytical units for assembly-state-resolved biomolecular quantification.

## MATERIALS AND METHODS

### Materials

Hen egg-white lysozyme (HEWL; cat. no. 10837059001), pepsin from porcine gastric mucosa (cat. no. P7000), Thioflavin T (ThT; cat. no. T3516), and the Avanti Mini-Extruder with holder/heating block were purchased from Sigma-Aldrich (USA). Halt™ protease inhibitor single-use cocktail (100×) was purchased from Thermo Fisher Scientific (USA). Track-etched polycarbonate (PC) membrane filters with a nominal pore size of 10 μm (Isopore, cat. no. TCTP01300) were purchased from Merck Millipore (USA). HEWL fibrils were mechanically truncated using 200 nm polycarbonate track-etched membranes (Whatman Nuclepore, Cytiva, cat. no. 10417004) during extrusion. Polydimethylsiloxane (PDMS) elastomer base and curing agent (Sylgard 184) were purchased from Dow (USA). Hydrochloric acid (HCl, 37%; Sigma-Aldrich, cat. no. 258148) and sodium hydroxide pellets (NaOH; Sigma-Aldrich, cat. no. 1064980500) were used to prepare 1 M HCl and 0.1 M NaOH stock solutions for pH adjustment. All experiments were performed using deionized water (DW) unless otherwise stated.

### APQ Chip Fabrication

The APQ chip was fabricated using a standard soft-lithography PDMS protocol.^58^ PDMS prepolymer was mixed with curing agent at a 10:1 (w/w) ratio, degassed under vacuum to remove air bubbles, and cured at 60 °C for 12 h. Two 1 mm-thick PDMS slabs (20 × 20 mm²) with a central circular opening of 10 mm diameter were prepared using a 10 mm biopsy punch. A track-etched PC membrane filter with a nominal pore size of 10 μm was sandwiched between the two PDMS slabs and aligned over the circular openings. The membrane was secured against each PDMS slab using double-sided paper tape with a thickness of 70 μm to prevent sample bypass around the membrane edges. The completed APQ chips were used immediately or stored at room temperature in a sealed container until use.

### Polycarbonate Membrane Pore Characterization

The pore-size distribution of the PC membrane (Figure 1b) was characterized by fluorescence imaging of pores filled with ThT solution. Images were acquired using an upright fluorescence microscope (Eclipse Ni, Nikon, Tokyo, Japan) equipped with a ThT-compatible filter cube and a 10× objective (1.0× tube lens) at an exposure time of 500 ms. Pore boundaries and diameters were measured from the acquired images using NIS-Elements BR software (Nikon, Tokyo, Japan). Pore-size statistics were aggregated from *n* = 4 independent fields of view.

### Preparation of HEWL Amyloid Fibrils

Deionized water was acidified to pH 2 using 1 M HCl. HEWL was dissolved at 2 wt% in pH 2 DW and filtered through a 0.2 μm polyvinylidene difluoride (PVDF) membrane syringe filter to remove preformed aggregates. The filtered solution was transferred to sealed glass vials and incubated in an oil bath at 65 °C for 3 days without agitation, following established protocols for HEWL fibril formation.^34, 59^ The resulting fibril stock was stored at 4 °C until use and diluted in pH 2 DW to the indicated working concentrations.

### t-ANF Formation via Extrusion

Truncated amyloid nanofibrils (t-ANFs) were prepared by mechanical shearing of the HEWL fibril stock (2 wt%) using an Avanti Mini-Extruder equipped with 1,000-μL syringes. The fibril solution was passed through a 200 nm PC membrane filter for 100 extrusion cycles at room temperature, following our previously reported protocol.^29^ Prior to extrusion, the membranes and filter supports were pre-wetted with DW to ensure consistent flow and reduce friction. The resulting t-ANF stock was stored at 4 °C and diluted in pH 2 DW to the working concentrations used in each experiment.

### Preparation of the HEWL Monomer and Oligomer Samples

Monomer samples were prepared by dissolving HEWL at 2 wt% in pH 2 DW and filtering the solution through a 0.2 μm PVDF membrane syringe filter immediately before use to remove preformed aggregates. Oligomer-rich HEWL samples were prepared using an axial-rotation–based mechanochemical protocol reported previously.^48^ Briefly, HEWL solutions were subjected to controlled axial rotation under conditions that favor soluble oligomeric assembly while suppressing mature fibril formation. Detailed preparation conditions and characterization are described in the cited report.^48^ Oligomer aliquots were used immediately or stored at 4 °C until use in APQ experiments.

### pH-Dependent t-ANF Aggregation Experiments

For pH-dependent aggregation experiments (Figure 4), t-ANF stock was diluted in DW to a working concentration of 0.1 wt%. The pH of each aliquot was independently adjusted to 9, 10, 11, or 12 using 0.1 M NaOH. The pH of each sample was verified using a calibrated benchtop pH meter (Orion Star A211, Thermo Fisher Scientific, USA). Each pH-adjusted t-ANF sample was loaded onto the APQ chip and processed under the standard vacuum-assisted ThT staining and imaging conditions described below. Aliquots from the same pH-adjusted samples were used to prepare AFM specimens (Figure 4e).

### Pepsin-Induced Amplification of t-ANFs

For pepsin-induced amplification experiments (Figure 5), t-ANF solutions (0.1 wt% in pH 2 DW) were mixed with pepsin to a final pepsin concentration of 2 μg/mL and incubated in a ThermoMixer C (Eppendorf, Germany) at 37 °C for 3 h without agitation. Four conditions were prepared in parallel: (i) t-ANFs alone (control), (ii) t-ANFs + pepsin, (iii) t-ANFs + pepsin + Halt™ protease inhibitor cocktail (final 1×; the inhibitor was added to the t-ANF solution before pepsin addition), and (iv) t-ANFs + heat-denatured pepsin. Heat-denatured pepsin was prepared by incubating the pepsin stock at 100 °C for 1 h and cooling it to room temperature before mixing with t-ANFs. After the 3-h incubation, samples were applied directly to the APQ chip. AFM specimens (Figure 5c) were prepared from aliquots of the t-ANF + pepsin condition using the same incubated stock.

### Monomer- and Oligomer-Induced t-ANF Amplification and pH-Induced Aggregation Experiments

For Stage 1 amplification experiments (Figure 6b,c), three conditions were prepared in parallel and incubated under matched conditions for 3 h at 60 °C in a ThermoMixer C (Eppendorf, Germany) without agitation: (i) t-ANFs alone (control), (ii) t-ANFs co-incubated with HEWL monomers, and (iii) t-ANFs co-incubated with HEWL oligomers. The final t-ANF concentration was 0.05 wt%, and the final monomer or oligomer concentration was 0.05 wt%, corresponding to a 1:1 (w/w) t-ANF-to-precursor ratio in pH 2 DW. After incubation, an aliquot of each sample was applied directly to the APQ chip and processed under the standard vacuum-assisted ThT staining and imaging conditions described below. For Stage 2 aggregation experiments (Figure 6d,e), separate aliquots of the same Stage 1 samples were adjusted to pH 10 using 0.1 M NaOH, gently mixed, loaded onto fresh APQ chips, and processed under the same standard staining and imaging conditions. The pH of each adjusted sample was verified using a calibrated benchtop pH meter (Orion Star A211, Thermo Fisher Scientific, USA).

### Vacuum-Assisted ThT Staining and APQ Measurements

Sample solutions were mixed with ThT to a final ThT concentration of 10 μM. An 80-μL aliquot of the mixed solution was dispensed onto the APQ chip, which was then placed in a vacuum desiccator. The desiccator was evacuated to −0.85 bar using a Gast vacuum pump (Gast Manufacturing, USA) and held under vacuum for 10 min, allowing the t-ANF/ThT solution to be drawn into the membrane pores.^30^ After infiltration, the chip was removed from the desiccator and imaged immediately. For optimization experiments (Figure 2), the infiltration time was varied (0, 1, 5, and 10 min) at a fixed ThT concentration of 10 μM, and the ThT concentration was varied (0, 1, 10, and 100 μM) at a fixed infiltration time of 10 min. All other experiments used the optimized standard condition of 10-min infiltration with 10 μM ThT.

### Fluorescence Imaging

Fluorescence images of the APQ chip were acquired using an upright fluorescence microscope (Eclipse Ni, Nikon, Tokyo, Japan) equipped with a 10× objective and a ThT-compatible filter set (excitation 450 nm; emission 490 nm). Images were captured using a DS-Ri2 camera with an integration time of 500 ms. To obtain representative pore-level distributions, five different regions of each membrane were imaged. Raw 16-bit grayscale images were saved without rescaling for downstream analysis.

### Pore-Based Intensity Analysis

Pore-level image analysis was performed using a custom MATLAB script provided as Supporting Information. Raw fluorescence images were analyzed without intensity rescaling. For each image, one representative pore point and one background point were manually selected to define an image-specific adaptive threshold:

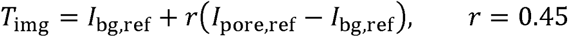

where *I*_pore,ref_ and *I*_bg,ref_ are the grayscale intensities at the selected pore and background reference points, respectively. Pixels above *T*_img_ were binarized, and connected components smaller than 20 pixels were removed using the bwareaopen function. Remaining connected components were labeled using bwlabel, and falsely detected objects were manually removed by selecting their centroid positions during a curation step. For each retained pore *i*, regionprops was used to extract the pore area *A_i_*, mean grayscale intensity *Ī_i_*, and centroid. The integrated pore intensity was calculated as:

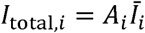

and exported as a comma-separated values file for downstream analysis. The selected background point was used only for adaptive thresholding and was not subtracted from the reported integrated pore intensity. Pore-level values were used to generate scatter plots, frequency histograms, heatmaps, and concentration-response curves.

### Bulk ThT Fluorescence Assay

For complementary bulk-level ThT measurements following standard protocols,^14^ samples were mixed with ThT to a final concentration of 10 μM, and 100-μL aliquots were transferred to black 96-well plates. Fluorescence intensity was measured using a microplate reader (Synergy H1, BioTek, USA) in top-read mode with excitation and emission wavelengths set to 445 and 485 nm, respectively. All measurements were performed with at least three technical replicates per condition.

### Atomic Force Microscopy

For AFM sample preparation, 10-μL aliquots of each sample were deposited onto freshly cleaved mica discs (Highest Grade V1, Ted Pella, USA) and incubated for 5 min at room temperature. The mica surface was then rinsed twice with 100 μL of DW and dried under a stream of N_ gas. AFM imaging was performed in tapping mode in air using a Multimode 8 system (Bruker, USA) with PR-T300 cantilevers (Probes; nominal spring constant, 40 N/m). Images were acquired at a scan rate of 0.6 Hz with a pixel resolution of ≤4 nm/pixel. AFM images were processed by plane-flattening and line-correction and analyzed using MountainsSPIP software (Digital Surf, France).

### Statistical Analysis

Data are presented as mean ± standard deviation unless otherwise noted. For APQ measurements, *N* denotes the number of analyzed pores. Statistical comparisons for bulk ThT fluorescence assays were performed using one-way ANOVA followed by Tukey’s post hoc test or unpaired two-tailed Student’s t-test, as appropriate. Significance levels were defined as ns, *p* ≥ 0.05; *\*p* < 0.05; *\*\*p* < 0.01; *\*\*\*p* < 0.001; *\*\*\*\*p* < 0.0001.

## ASSOCIATED CONTENT

### Supporting Information

Z-factor analysis of pore-level discrimination, concentration-dependent line-scan profiles, chip-to-chip reproducibility, bulk ThT fluorescence controls, AFM-based fibril size analysis, temperature-dependence measurements, AFM topography of monomer- and oligomer-processed t-ANFs, and pH-adjusted amyloid assemblies (PDF). MATLAB script for APQ pore segmentation and pore-level intensity extraction (M file).

## AUTHOR INFORMATION

### Author Contributions

The manuscript was written through the contributions of all authors. All authors have given approval to the final version of the manuscript.

Seokbeom Roh: Conceptualization, Investigation, Data curation, Methodology, Formal analysis, Visualization, Writing – original draft.

Sechan Han: Investigation, Data curation, Methodology, Formal analysis, Visualization, Writing – original draft.

Minwoo Bae: Investigation, Methodology, Validation.

Eunji Song: Investigation, Methodology, Validation.

Taeha Lee: Formal analysis, Validation.

Dain Kang: Investigation, Formal analysis. Da Yeon Cheong: Investigation, Validation.

Hyungbeen Lee: Investigation, Validation.

Soochi Kim: Investigation, Validation.

Gyudo Lee: Conceptualization, Investigation, Supervision, Funding acquisition, Project administration, Writing – review & editing.

### Notes

The authors declare no competing financial interest.

## ACKNOWLEDGMENT

This research was supported by the National Research Foundation of Korea (NRF) (RS-2025-16068427), Ministry of Health and Welfare (KH140292), Ministry of Science and ICT (IITP-2026-RS-2023-00258971), and a Korea University Grant.

## ABBREVIATIONS

APQ: amyloid pore quantification
DW: deionized water
HEWL: hen egg-white lysozyme
pI: isoelectric point
t-ANF: truncated amyloid nanofibril
ThT: thioflavin T.

