## Supplementary Information for "Pore-Resolved High-Throughput Quantification of Amyloid Aggregation and Amplification"

**S1.** **Z-factor analysis of pore-level discrimination between conventional fibrils and t-ANFs**

The pore-level separation between conventional fibrils and t-ANFs was evaluated using the Z-factor:

$$Z^{'}=1-\frac{3(\sigma_{\mathrm{fibril}}+\sigma_{t-\mathrm{ANF}})}{\mid\mu_{t-\mathrm{ANF}}-\mu_{\mathrm{fibril}}\mid}$$

where $\mu$ and $\sigma$ represent the mean and standard deviation of the single-pore ThT intensity distributions, respectively. The calculated Z-factor was 0.614, indicating a robust assay window between conventional fibrils and t-ANFs. This result confirms that extrusion-mediated truncation improves not only the average pore intensity but also the statistical separability of the pore-resolved readout.

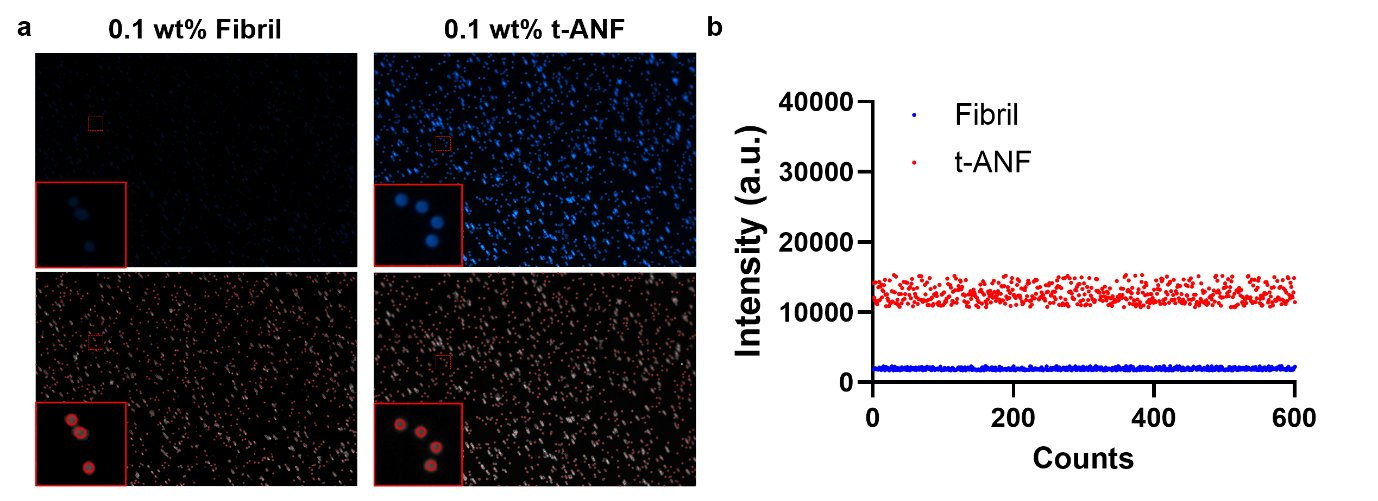

**Figure S1.** Pore-resolved fluorescence comparison between conventional amyloid fibrils and truncated amyloid nanofibrils (t-ANFs) at matched concentration (0.1 wt%). (a) Fluorescence images of pore fields stained with ThT. Top row: ThT signal channel; bottom row: detected pore boundaries (red circles) overlaid on a darkened reference image. The 0.1 wt% conventional fibril sample (left) yields sparse and dim pore signals due to limited pore accessibility of long fibrils, whereas the 0.1 wt% t-ANF sample (right) yields uniformly bright pores. Insets (red boxes) show magnified views of representative pores. (b) Single-pore ThT intensity scatter plot (N = 600 pores per condition). t-ANF (red) yields a mean intensity of ~13,000 a.u., approximately one order of magnitude higher than conventional fibrils (blue, ~2,000 a.u.). These results demonstrate that mechanical truncation by extrusion is essential for quantitative pore-resolved analysis.

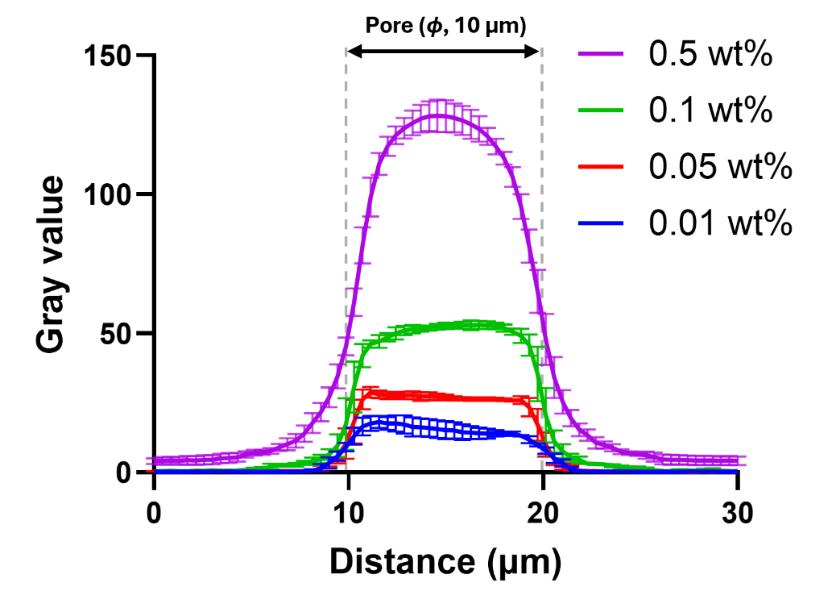

**Figure S2.** Concentration-dependent line-scan gray-value profiles across single pores filled with t-ANF at 0.01, 0.05, 0.1, and 0.5 wt%. Profiles are averaged across 5 pores per condition; error bars represent standard deviation. The 10 μm pore region (φ) is delimited by gray dashed lines. The flat-topped distributions indicate uniform fluorescence across each pore, and the peak gray value scales monotonically with t-ANF concentration. The data support the use of integrated pore fluorescence intensity as the quantitative readout in Figure 3.

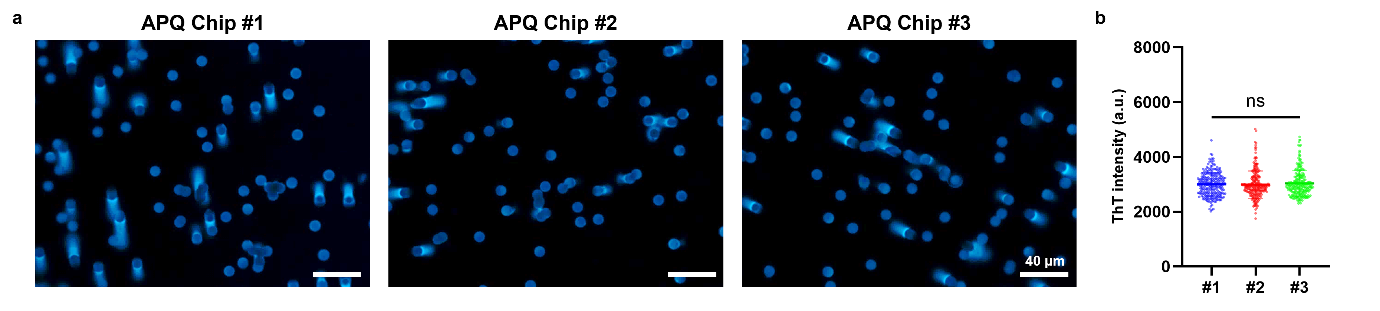

**Figure S3.** Reproducibility of APQ chip-based t-ANF quantification. (a) Representative fluorescence images of three independent APQ chips loaded with the same 0.1 wt% t-ANF sample. Scale bars: 40 μm. (b) Pore-level ThT intensity distributions obtained from each APQ chip, showing comparable pore-level intensity distributions among independent chips. Data indicate reproducible chip-level performance of the APQ readout under the reference t-ANF condition.

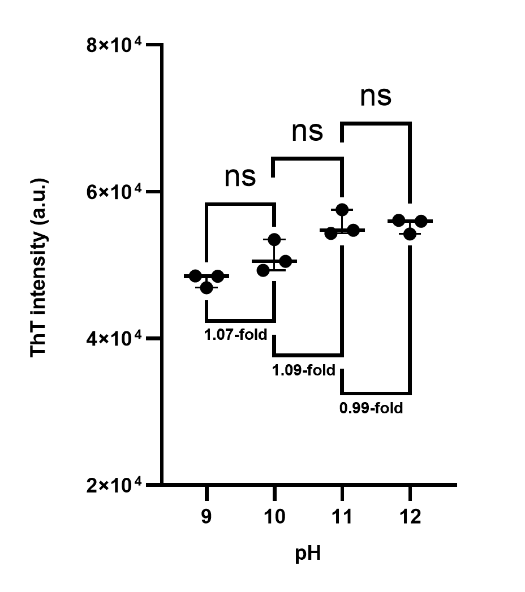

**Figure S4.** Bulk Thioflavin T (ThT) fluorescence intensity of t-ANF samples incubated at pH 9, 10, 11, and 12, measured in solution by microplate reader (excitation 445 nm / emission 485 nm; n = 3 per condition; bars indicate mean ± SD). Pairwise fold differences between adjacent pH values are 1.07, 1.09, and 0.99, with no statistically significant differences detected (one-way ANOVA, ns). In contrast to the APQ measurement (Figure 4), bulk ThT cannot resolve pH 11–specific aggregation behavior because cross-β content is preserved regardless of aggregate size. This comparison highlights the size-discriminating advantage of the pore-based assay.

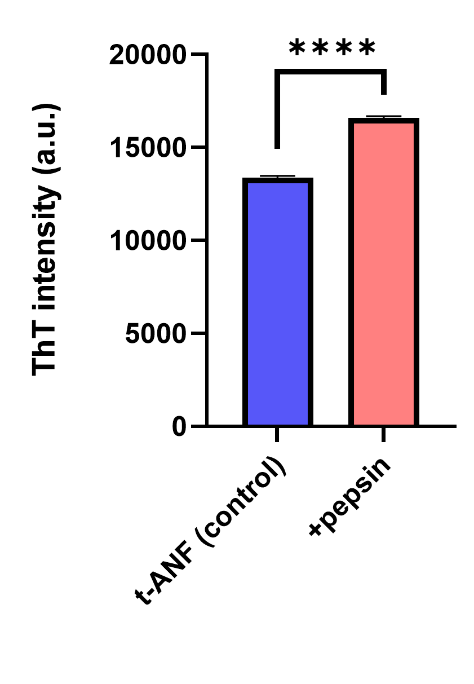

**Figure S5.** Bulk ThT fluorescence intensity of t-ANF (control) and t-ANF + pepsin samples (37 °C, 3 h incubation; n = 3 each), measured in solution by microplate reader. Pepsin treatment yields a statistically significant increase in ThT signal (****, p < 0.0001, two-tailed unpaired t-test). Although bulk ThT detects an average increase in cross-β content, it cannot resolve fibril fragmentation, secondary elongation, or the size-dependent pore-passage behavior captured by the APQ platform (Figure 5).

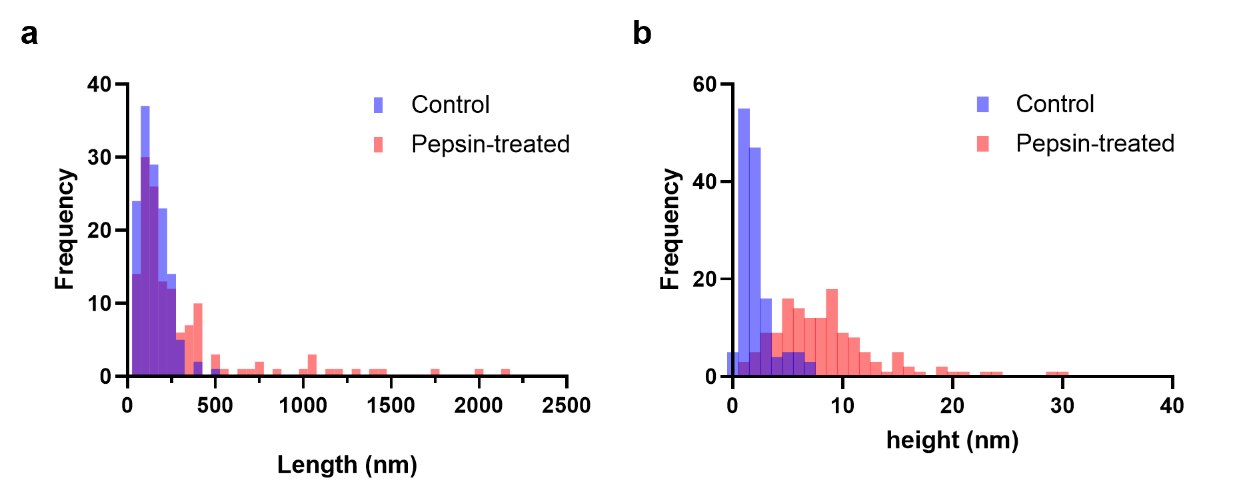

**Figure S6.** AFM-based size analysis of t-ANF before and after pepsin treatment. (a) Length distribution histograms of control t-ANF (blue, n ≥ 100 fibrils) and pepsin-treated t-ANF (pink, n ≥ 100 fibrils). Pepsin treatment produces both a sub-population of shortened fragments (<200 nm) and a long tail of elongated species extending up to ~2,000 nm. (b) Height distribution histograms for the same samples. The control distribution is narrow and centered near ~2 nm, characteristic of single t-ANFs, whereas the pepsin-treated distribution shifts markedly to higher values (broad range up to ~30 nm), consistent with bundling of newly elongated species. The bimodal length and elevated height profiles support a fragmentation-coupled secondary elongation mechanism (Figure 5).

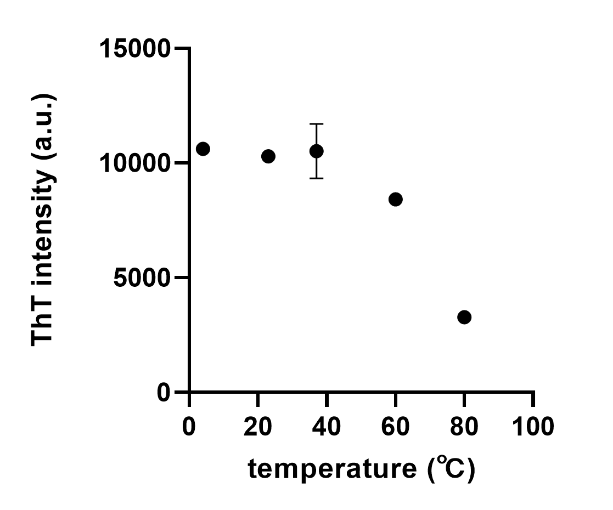

**Figure S7.** Temperature dependence of bulk ThT fluorescence for t-ANF, measured after 3 h incubation at 4, 23, 37, 60, and 80 °C (n = 3 per condition; error bars indicate SD where larger than the marker). The ThT signal remains essentially constant from 4 to 37 °C (~10,000 a.u.), decreases modestly at 60 °C (~8,400 a.u.), and drops sharply at 80 °C (~3,200 a.u.), consistent with progressive thermal disassembly of fibrils. The 60 °C condition was selected for the monomer/oligomer amplification experiments (Figure 6) as it accelerates amplification kinetics while preserving the structural integrity of t-ANFs.

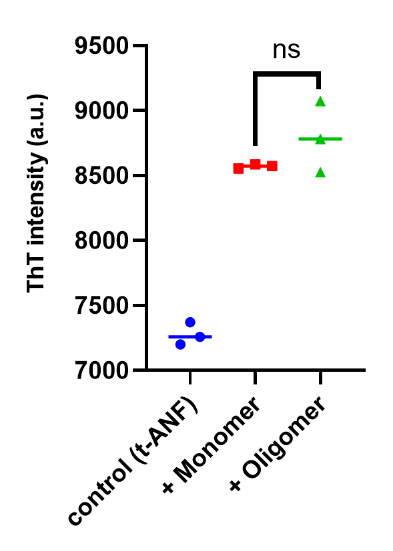

**Figure S8.** Bulk ThT fluorescence intensity of t-ANF (control), t-ANF + monomer, and t-ANF + oligomer samples after 3 h incubation at 60 °C (n = 3 per condition; bars indicate mean). Although both monomer and oligomer addition increase the bulk signal compared with the control (~7,300 a.u.), the +monomer (~8,570 a.u.) and +oligomer (~8,790 a.u.) conditions are not statistically distinguishable (ns, two-tailed unpaired t-test). This contrasts with the clear three-way discrimination achieved by the APQ platform (Figure 6c), demonstrating that bulk ThT cannot distinguish secondary nucleation from elongation pathways.

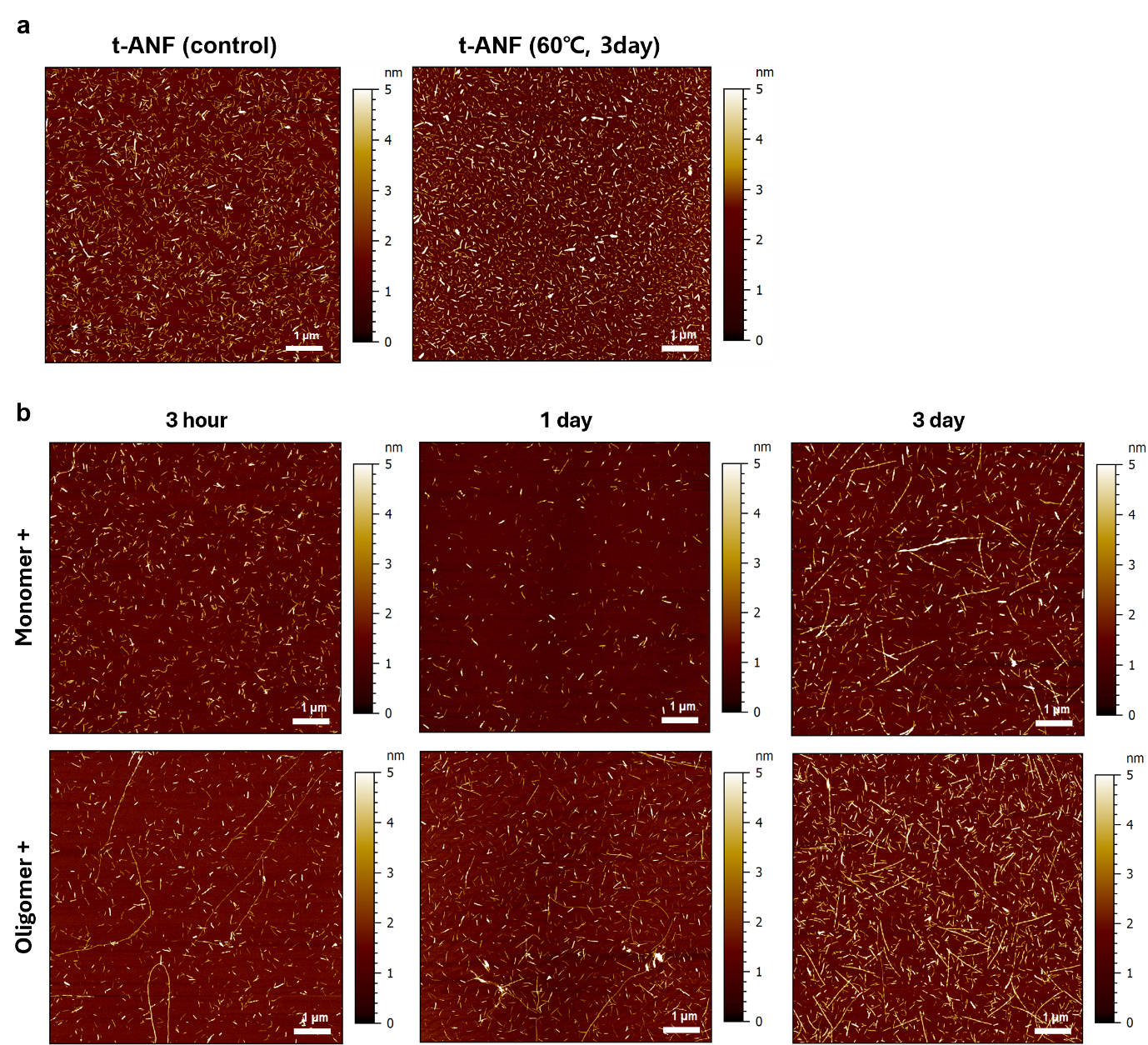

**Figure S9.** AFM topography of t-ANF morphological evolution under different conditions. (a) AFM images of control t-ANF (left) and t-ANF after 3 days of incubation at 60 °C without seeds (right). Fibril morphology and density are preserved, confirming that the APQ signal changes observed in Figure 6 originate from interactions with added monomer or oligomer rather than from autonomous growth. (b) Time-resolved AFM images of t-ANF + monomer (top row) and t-ANF + oligomer (bottom row) at 3 h, 1 day, and 3 day incubation. Monomer-seeded samples show only modest elongation over time, whereas oligomer-seeded samples display markedly elongated, network-like fibrils as early as 3 h, with extensive entanglement by 3 day. These observations support the enhanced elongation pathway proposed in Figure 6a. All images use a height color scale of 0–5 nm; scale bars: 1 μm.

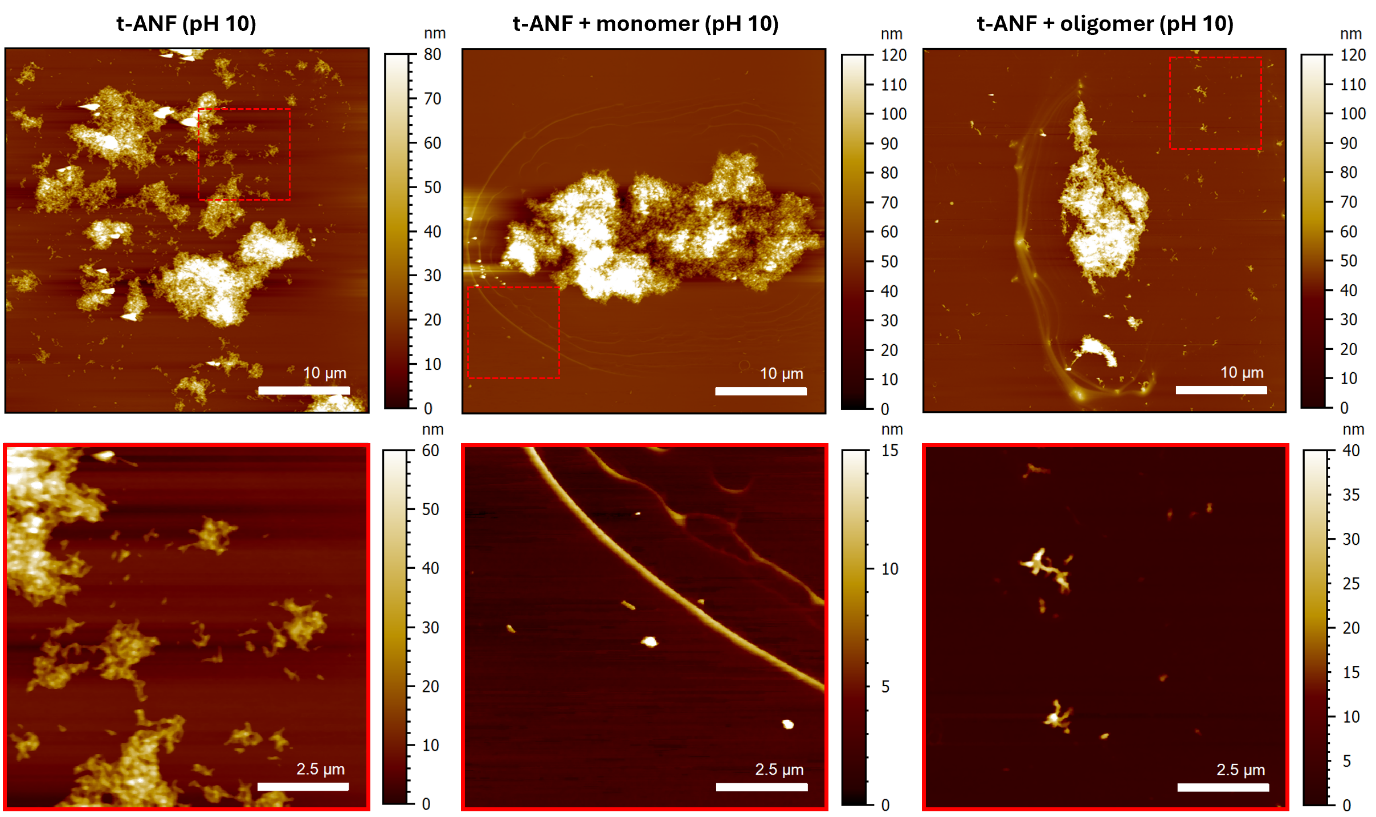

**Figure S10.** AFM characterization of pH-adjusted amyloid assemblies after short-term incubation. Representative AFM height images of t-ANF alone, t-ANF incubated with monomers, and t-ANF incubated with oligomers after incubation at 60 °C for 3 h and subsequent adjustment to pH 10. The lower panels show magnified AFM images of the regions indicated by the red dashed boxes in the corresponding upper panels. Scale bars: 10 μm in the upper panels and 2.5 μm in the lower panels.
